# Striatal seeding of protofibrillar alpha-synuclein causes cortical hyperreactivity in behaving mice

**DOI:** 10.1101/2020.09.28.314526

**Authors:** Sonja Blumenstock, Fanfan Sun, Petar Marinković, Carmelo Sgobio, Sabine Liebscher, Jochen Herms

## Abstract

Alpha-synucleinopathies are characterized by self-aggregation of the protein alpha-synuclein (a-syn), causing alterations on the molecular and cellular level. To unravel the impact of transneuronal spreading and templated misfolding of a-syn on the microcircuitry of remotely connected brain areas, we investigated cortical neuron function in awake mice 9 months after a single intrastriatal injection of a-syn preformed fibrils (PFFs), using *in vivo* two-photon calcium imaging. We found altered function of layer 2/3 cortical neurons in somatosensory cortex (S1) of PFF-inoculated mice, as witnessed by an enhanced response to whisking and increased synchrony, accompanied by a decrease in baseline Ca^2+^ levels. Stereological analyses revealed a reduction in GAD67-positive inhibitory cells in S1 in PFF-injected brains. These findings point to a disturbed excitation/inhibition balance as an important pathomechanism in alpha-synucleinopathies and demonstrate a clear association between the spread of toxic proteins and the initiation of altered neuronal function in remotely connected areas.

## Introduction

Several neurodegenerative disorders share a pathological mechanism, by which the misfolding of a native monomeric protein into a range of intermediate oligomeric structures is followed by the deposition of amyloid fibrillary protein inclusions over a prolonged course of years to decades. Parkinson’s disease (PD) and other alpha-synucleinopathies are characterized by the misfolding, self-aggregation and deposition of the native protein a-synuclein (a-syn), causing various dysfunctions in e.g. ER-to-Golgi trafficking, cytoskeleton dynamics, protein degradation, synaptic vesicle cycle and calcium homeostasis (Reznichenko et al., 2012; Wong and Krainc, 2017). Extensive evidence supports the concept of “molecular” and “structural” templating, referring to conformational changes of the a-syn protein, subsequent accumulation and its spread along anatomically connected structures (Li et al., 2008; Uchihara and Giasson, 2016), which can be initiated by various molecular forms of a-syn such as monomers, oligomers or fibrils. We and others have employed the infusion of preformed, native, recombinant a-syn fibrils (PFFs) to initiate its seeding and spreading as a model to study a-synucleinopathies *in vivo* (Blumenstock et al., 2017; Luk et al., 2012; Masuda-Suzukake et al., 2013; Osterberg et al., 2015; Volpicelli-Daley et al., 2011). When applied acutely in cell culture, PFFs have recently been shown to compromised neuronal excitability and activity, to trigger intracellular a-syn pathology and to cause a rapid loss of spines (Wu et al., 2019). In our previous work, we already observed spine loss on apical dendrites of layer 5 cortical neurons months after striatal a-syn seeding (Blumenstock et al., 2017).

In the current study, we thus investigated the impact of striatal a-syn seeding on neuronal function in the upstream somatosensory cortex using two-photon *in vivo* calcium imaging in awake, behaving mice. We recorded neuronal activity in the somatosensory cortex 9 months after the unilateral infusion of PFFs into the dorsal striatum, which triggered the development of hyperphosphorylated, fibrillary and Lewy-neurite like a-syn inclusions in the cortex. Compared to controls, PFF-seeded mice exhibited an increase in the frequency as well as in the amplitude of spontaneous calcium transients in cortical layer 2/3. Most importantly, we observed a large increase in neuronal responsiveness (hyperreactivity) to whisking, accompanied by a higher pairwise correlation. Mechanistically, we could attribute these changes to a loss of GABAergic interneurons, causing a compromised inhibition within the microcircuitry.

## Results

### Injection of a-syn preformed fibrils induces formation of Lewy-neurite like aggregates in wild-type mice

As we have shown in a previous study and also demonstrate here, the injection of a-syn PFFs into the dorsal striatum triggers the formation of dense intracellular Lewy-neurite like aggregates in remotely connected areas in the brain. Here, we show that intracellular phosphorylated a-syn aggregates can be observed in somatosensory cortex 9 months after the injection of PFFs.

Aggregates can be observed across all cortical layers, but shows a higher density in infragranular layers, which contains extensive projections to the dorsal striatum. Importantly, aggregates are also present in the layers 2/3 of the somatosensory cortex (Suppl. Fig. S1).

### Hyperreactivity in somatosensory cortex in a-syn PFF injected mice

To investigate functional consequences of a-syn templated misfolding in the intact mouse cortex, we performed *in vivo* two-photon calcium imaging in awake, head-fixed mice 9 months after seeding with PFFs (Fig. 1A-C). We observed a pronounced increase in responsiveness to individual whisking events both at the population level, as well as on the level of individual neurons (Fig. 1D-F). More specifically, the average response of active neurons to individual onsets of whisking was significantly increased in a-syn mice (Fig. 1E, area under the curve from 0.5 – 3 seconds after whisking onset (grey area); *P* < 10^−19^, student’s t test). Furthermore, the fraction of neurons in each individual experiment, which were responsive to whisking (i.e. neurons that display a significant increase in ΔF/F within 0.5-1.5 seconds after whisking onset compared to their activity within 0.5 seconds prior to whisking onset) was also significantly increased in a-syn (*P* = 0.003, ranksum test, control n = 32 experiments, a-syn n = 32 experiments, Fig. 1F). We also observed a right shift in the distribution of transient frequencies and amplitudes during quiescence (spontaneous activity, frequency *P* = 0.016, Kolmogorov-Smirnov (KS) test; amplitude *P* = 0.0019, control n = 1561 ROIs, a-syn n = 1534 ROIs; Suppl Fig. S2A,B) and during whisking-epochs (frequency *P* = 0.011; amplitude *P* = 0.02, KS test; control n = 1561 ROIs, a-syn n = 1534 ROIs; Suppl Fig. S2A,B) in a-syn compared to control mice. The overall time spend whisking did not differ between control and a-syn mice (*P* = 0.45, ranksum test). The level of spontaneous activity correlated well with the whisking-associated activity levels in both control and a-syn mice (Fig. 1G, control R^2^= 0.63, y = 1.36x – 0.07; *P* < 0.0001; a-syn R^2^ = 0.63, y = 1.26x + 0.44, *P* < 0.0001). To characterize and quantify the relationship between spontaneous versus whisking-associated neuronal activity for each cell, we computed the angle of each data point in a plot displaying the whisking-associated versus spontaneous frequency (Fig. 1G). This analysis revealed that the distribution of angles was significantly different in a-syn mice (*P* < 10^−4^, KS test, Fig. 1H), with a much larger fraction of neurons favoring activity during whisking over spontaneous activity. Neuronal activity did also not differ between control and a-syn mice during anesthesia (Suppl. Fig 2C,D). In addition, we addressed the question whether baseline calcium levels might also be affected in a-syn PFF-seeded mice. To this end, we took advantage of the stoichiometric expression of both fluorophores mRuby2 and GCaMP6s through the AAV2/1.hSyn1.mRuby2.GSG.P2A.GCaMP6s. We thus computed the ratio between the baseline values of the green and red channel (G/R ratio). On average neurons in a-syn PFF-seeded mice had a slightly, yet significantly lower ratio compared to control mice (control: 0.139 (0.138 – 0.141), a-syn: 0.132 (0.129 – 0.134), data are median + 95% CI of the median, *P* < 0.001, ranksum test. Suppl. Fig. S2), indicating that the baseline calcium content is even lower in neurons in a-syn PFF-seeded mice. Pairwise neuronal correlations during stationary epochs were not affected (*P* = 0.43, KS test, Fig. 2A-C), while whisking-associated neuronal activity appeared more correlated (control vs a-syn *P* = 0.026, control vs shuffled control *P* < 0.0001, a-syn vs shuffled a-syn *P* < 0.0001, KS test, Fig. 2D) upon PFF injection.

**Figure 1.**
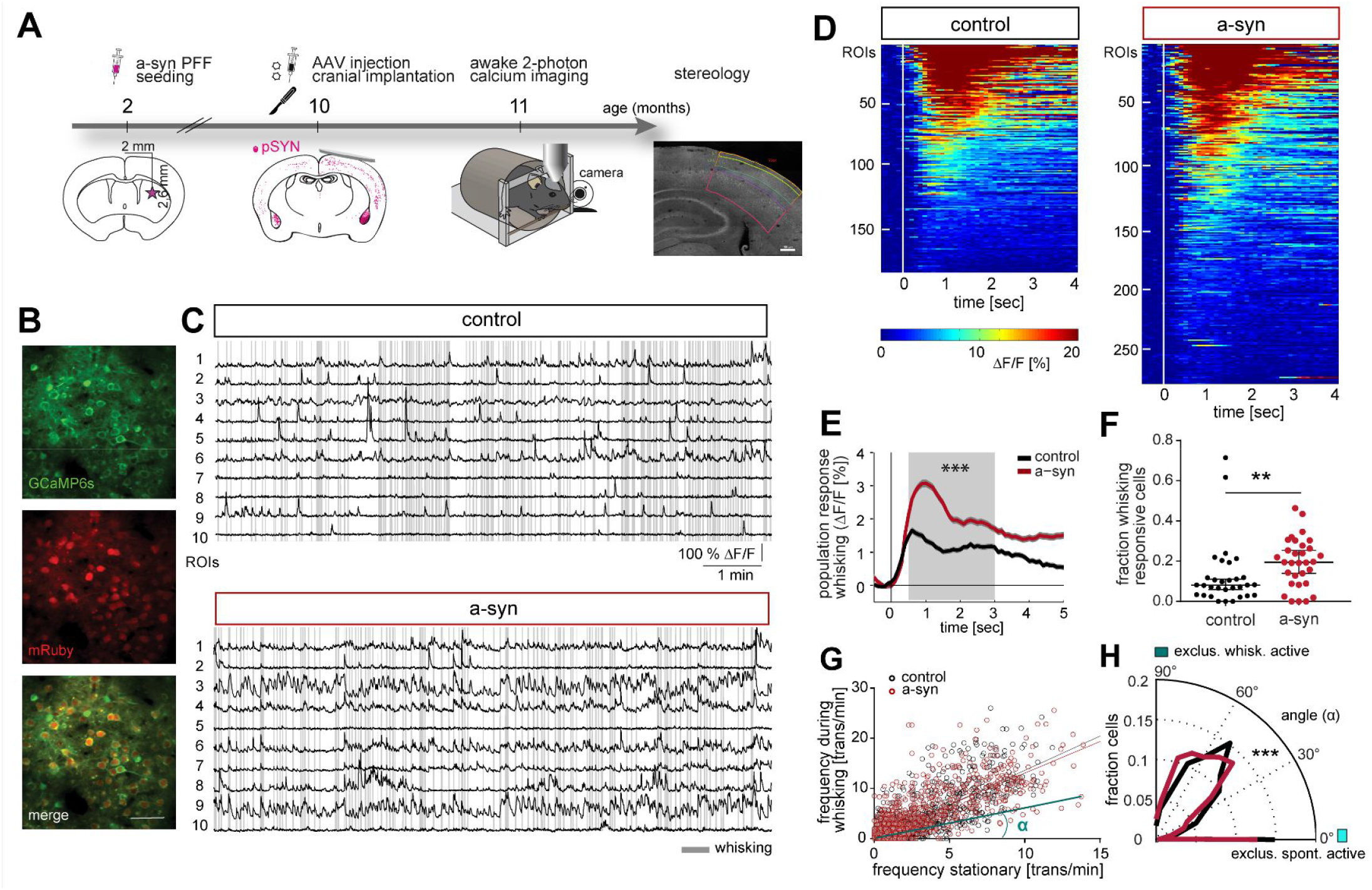
Neuronal hyperreactivity in S1 upon striatal injection of a-syn PFFs. **(A)** Timeline of experiments. Mice received a striatal injection of PFFs at the age of 2 months, followed by the injection of AAV2/1.hSyn1.mRuby2.P2A.GCaMP6s into the somatosensory cortex (S1) and the implantation of a cranial window 8 months later when PFFs are globally present (assessed by immunofluorescence of phospho-synuclein (pSYN)). One month later *in vivo* imaging experiments were performed, after which mice were sacrificed and stereology was conducted. **(B)** Representative example of a field of view (FOV). **(C)** Calcium traces of individual regions of interest (ROIs) are shown for control and a-syn mice and referenced by whisking activity (gray lines). **(D)** Heat maps depicting the average neural response to whisking onset (white line, normalized to the average activity within 1 sec before whisking onset) for whisking responsive cells (control: 187 of 1561 neurons, a-syn: 276 of 1534 neurons). **(E)** Population response of active neurons to whisking onset. **(F)** Fraction of whisking responsive neurons in each experiment. **(G)** Relationship of the stationary neuronal frequency and the activity associated with whisking in control (black) and a-syn mice (red). To compare state-specific neuronal activity levels, the angle α was computed for each neuron, as exemplified for one neuron (light blue line and angle). **(H)** The distribution of all angles is significantly different in a-syn mice, with more neurons favoring activity during whisking epochs (angle of 0° indicates neuronal activity exclusively during stationary (quiescent) epochs, while 90° would indicate exclusive whisking-associated neuronal activity). Data are mean +/- SEM. Scale bar in B is 50μm. ** *P* < 0.01, *** *P* < 0.001

**Figure 2.**
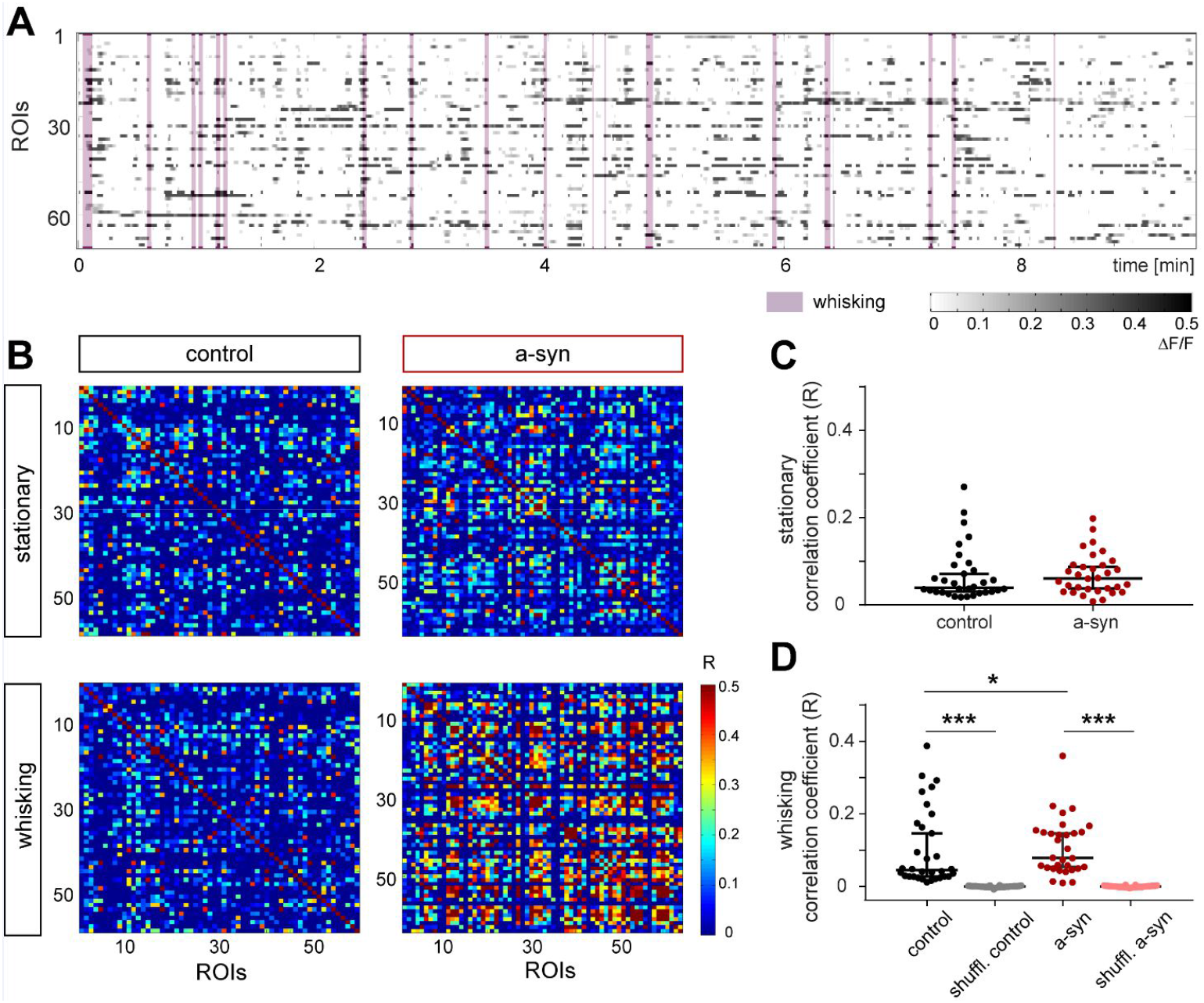
Striatal PFF seeding elevates pairwise neuronal correlations during whisking in S1. **(A)** Example raster plot depicting activity of each ROI within an FOV referenced by whisking (purple area). **(B)** Correlograms of pairwise correlations during stationary and whisking-associated epochs in a control and an a-syn mouse. **(C)** Average pairwise correlations of individual experiments did not differ, while **(D)** whisking-associated correlations were significantly increased in a-syn compared to control mice and each to shuffled data. Data are individual experiments superimposed by the median +/- 95% confidence interval (C,D). * *P* < 0.05, *** *P* < 0.0001.

**Figure 3.**
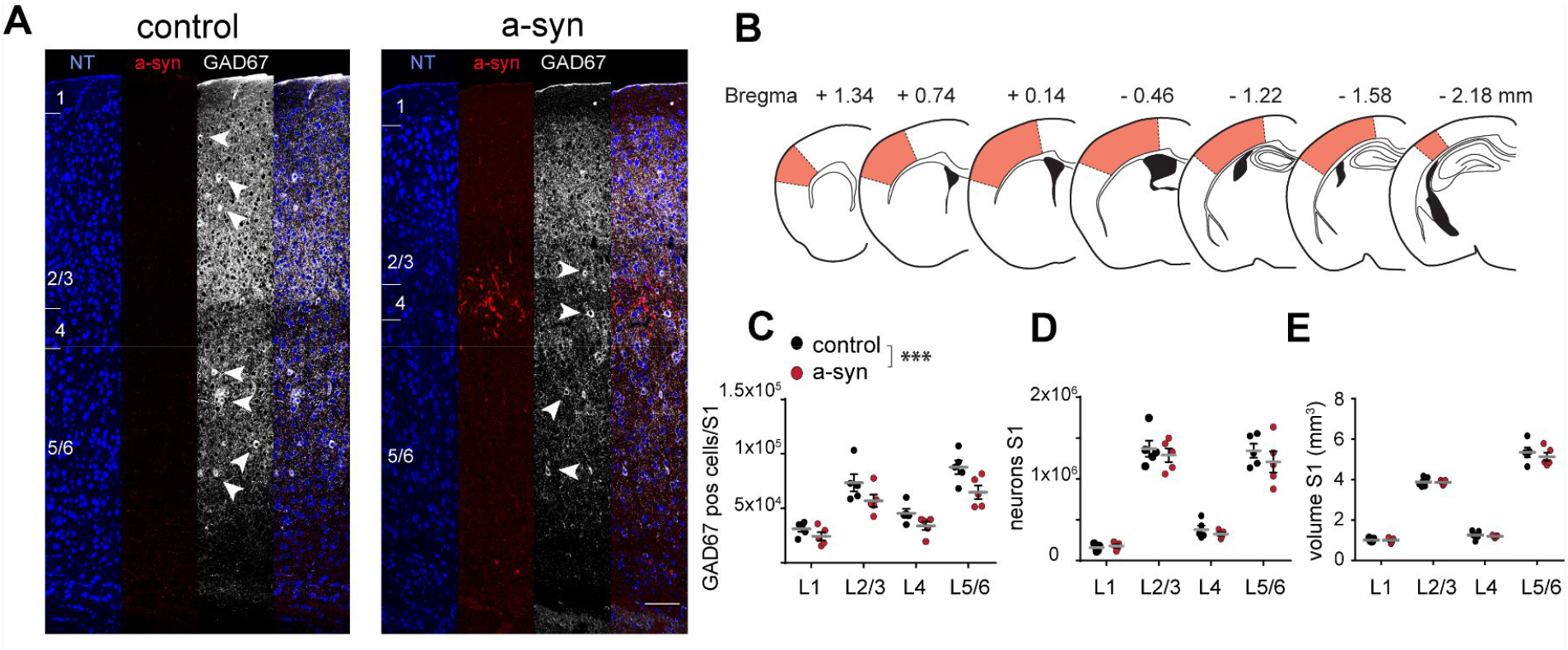
Stereology of somatosensory cortex reveals reduction of inhibitory neurons. (A) Representative examples of immunohistochemical stainings for neurotrace (NT), phospho-synuclein (a-syn, red) and GAD67 (white arrow heads mark GAD67 positive GABAergic neurons) in S1 of a control and an a-syn PFF-seeded mouse assessed 10 months after striatal seeding. (B) Brain sections used for stereology to assess overall number of neurons and of GABAergic cells across all cortical layers in the entire S1 area (orange area). (C) The number of GAD67 positive neurons was significantly reduced in a-syn mice, while neither (D) the total number of all neurons, nor (E) the cortical volume of S1 was affected. Data are mean +/- SEM in (C,D,E). Scale bar 100μm in (A). *** *P* < 0.0001.

### Stereological analysis in imaged brains reveals reduction in GABAergic interneuron numbers

We found a reduction in the number of GAD67-positive inhibitory cells (two-way ANOVA, effect of group F(1,32) = 14.69, *P* = 0.0006; effect of layer F(3,32) = 35.12, *P* < 0.0001), which was most prominently seen for layer 5/6 (*P* = 0.019; Bonferroni *post-hoc* test, all other layers n.s.). This reduction was specific the inhibitory neurons as the overall number of neurons was not affected (effect of group F(1,32) = 1.41, *P* = 0.24, effect of layer F(3,32) = 132.2, *P* < 0.0001) and also the cortical volume was not changed (effect of group F(1,32) = 0.78, *P* = 0.39, effect of layer F(3,32) = 530.3, *P* < 0.0001) by the injection of PFFs.

## Discussion

Since the identification of alpha-synuclein (a-syn) as the principal component of Lewy bodies and Lewy neurites (Spillantini et al., 1998; Spillantini et al., 1997), many studies have suggested its causative role in the pathogenesis of PD and Lewy Body Dementia. The intra- and extracellular concentration of a-syn as well as its protein conformation and degree of polymerization critically determine the onset of neurodegenerative processes (Henderson et al., 2019; Luk et al., 2012; Peelaerts et al., 2015; Peng et al., 2020).

Propagation and templated misfolding of a-syn in the brain is known to occur in both anterograde and retrograde direction, as well as through local release and diffusion. Growing evidence even suggests a critical role of the peripheral nervous system and gut a-syn in the onset of the pathologic cascade (Challis et al., 2020; Chen et al., 2018; Shrivastava et al., 2017; Wang et al., 2020).

We have previously shown that synapses in S1 are vulnerable to misfolded a-syn several months after striatal seeding. However, whether this propagation of a-syn aggregation is sufficient to alter neuronal function and if so, to what degree, remains unknown.

To address these questions, we recorded neuronal activity in awake, behaving mice using two-photon calcium imaging 9 months after a single intrastriatal injection of a-syn PFFs. Striatal seeding of PFFs caused neuritic as well as mature somatic inclusions across all cortical layers in S1, a region upstream of the inoculated striatum, and which had been previously shown to also bear hallmarks of human Lewy pathology, such as ubiquitination and amyloid conformation (Blumenstock et al., 2017; Osterberg et al., 2015).

Our *in vivo* imaging approach revealed a pronounced increase in neuronal activity in response to whisking (hyperreactivity) in a-syn PFF-injected mice. The observed elevated population response in S1 can be attributed to an increase in the fraction of whisking responsive cells and suggests an altered excitation/inhibition balance. Furthermore, stronger pairwise neuronal activity correlations indicate increased network synchrony. Importantly, the elevation in cortical activity did not extend to experiments conducted under anesthesia, indicating the importance of a behavioral stimulus (i.e. whisking) for the observed effect. Acute application of PFFs has recently been shown to reduce neuronal activity and cause spine loss (Wu et al., 2019). Our data, however, suggest that under chronic conditions *in vivo* PFF seeding can cause the opposite effect on the network level.

Mechanistically, we could attribute the observed hyperreactivity to a loss of ~25% of GAD67 positive interneurons in S1 in a-syn PFF-seeded mice, occurring after the long incubation period of 9 months. This pronounced drop in overall GABAergic neuron number is highly likely causing an inhibitory deficit within the local microcircuitry. Our results thus provide important mechanistic insight into the molecular impact of a-syn aggregates on neuronal subtypes and provides the first *in vivo* evidence that accumulation of wild-type a-syn causes functional aberrations of cortical neurons over a prolonged time. Earlier studies have demonstrated the disturbance of calcium homeostasis and an increase in excitatory transmission in response to pathological a-syn and other amyloidogenic proteins (Demuro et al., 2005; Hüls et al., 2011; Reznichenko et al., 2012). This earlier work argues for an elevation of intracellular Ca^2+^ based on a-syn oligomer-mediated pore formation in the postsynaptic membrane (Schmidt et al., 2012; Volles et al., 2001), enhancement of voltage-operated Ca^2+^ channel activity (Hettiarachchi et al., 2009) or inefficient membrane repolarization (Shrivastava et al., 2015). However, most of these reports are based on studies of cultured cells or artificial phospholipid membranes and rely on overexpression or acute application of high concentrations of a-syn. In the *in vivo* scenario, we here in fact observed a reduction of baseline Ca^2+^ levels. Interestingly, a recent publication is showing reduced cytosolic Ca^2+^ concentration at early stages of the degenerative process in cultured cells, while only at later stages Ca^2+^ levels increased (Betzer et al., 2018). It is important to remember that in both our seeded mice and in humans, the majority of cortical neurons are actually free of Lewy-neurite like aggregates and therefore are likely exposed to disease-mechanisms other than simply an overload of fibrillar a-syn. Nevertheless, the prolonged cellular state of progressive a-syn aggregation in a subset of neurons in our model more closely reflects the mostly sporadic human pathophysiology than transgenic mice overexpressing a-syn or cultured cells and tissues do.

Cortical network hyperactivity has been described in mouse models of AD and HD (Arnoux et al., 2018; Burgold et al., 2019; Busche et al., 2008; Liebscher et al., 2016; Marinković et al., 2019) and a recent study demonstrated spontaneous LFP elevations in the beta- and gamma-bands in the olfactory bulb and piriform cortex of mice observed *in vivo* after a-syn seeding (Kulkarni et al., 2020). Importantly, hyperactivation of sensorimotor cortical areas was also shown in electrophysiological and functional magnetic resonance imaging studies conducted in parkinsonian rats and patients and have been attributed to both motor and psychotic symptoms in synucleinopathies (Li et al., 2012; Nagahama et al., 2010; Yu et al., 2007). These data strongly emphasize the translational relevance of our findings also for human pathophysiology.

Importantly, our data now indicate that the observed hyperreactivity is based on compromised inhibition due to a loss of GABAergic interneurons. While the exact causal link between cortical network disturbance and the observed reduction in interneuron numbers as well as the molecular mechanisms underlying this cell-type specific vulnerability still need to be investigated in more detail, we can conclude from our findings that a disturbance of excitation/inhibition balance could be a main driver of cortical dysfunction in response to local a-syn aggregation in synucleinopathies.

Dysregulation of cortical activity in turn could through corticostriatal projections lead to profound changes in the downstream striatum. A disturbed balance in the activity of direct and indirect spiny projection neurons has long been associated with dyskinesia in PD (McGregor and Nelson, 2019), but the classical rate model of parkinsonism has been challenged (Parker et al., 2018). Functional studies at cellular resolution are therefore crucial for defining the exact relationships in parkinsonian networks.

In conclusion, we provide compelling evidence for a functional impact on remotely connected brain areas after fibrillary a-syn seeding, likely conveyed through the selective vulnerability of interneurons. Importantly, our data demonstrate that chronic effects of a-syn accumulation can strongly differ from acute effects, causing an opposing impact on the network level through celltype specific vulnerability. To the best of our knowledge, this is the first study to demonstrate changes in neuronal activity assessed 9 months after the seeding of a-syn preformed fibrils in awake, behaving wild-type mice.

It further sustains the hypothesis that elevated levels and spreading of a-syn alters the excitatory/inhibitory balance in cortical circuits, which is likely based on dendritic spine pathology underlying synucleinopathies. Future work investigating the causal relationship between the structure and activity of selected principal neuron and interneuron populations will be important to understand the cell-type specific vulnerability to a-syn pathology.

## Acknowledgements

This work was funded by the Deutsche Forschungsgemeinschaft (DFG, German Research Foundation) under Germany’s Excellence Strategy within the framework of the Munich Cluster for Systems Neurology - EXC 2145 SyNergy – ID 390857198 (SL, JH), EXC 1010 (SB), the DFG, Emmy Noether Programme (SL). We are particularly grateful to Pieter Goltstein for developing and sharing software for data analysis. We also thank Eric Grießinger, Sarah Hanselka, and Nadine Lachner for excellent technical support. We are grateful to the Genetically-Encoded Neuronal Indicator and Effector (GENIE) Project and the Janelia Farm Research Campus of the Howard Hughes Medical Institute, in particular to Vivek Jayaraman, Rex A. Kerr, Douglas S. Kim, Loren L. Looger, and Karel Svoboda for developing and distributing the genetically encoded calcium indicator GCaMP6s.

## Author Contributions

Conception and design of study (SB, JH), data acquisition (SB, FS), setup design / data acquisition assistance (PM) software development (SL), data analysis (SB, FS, CS, SL), data interpretation (SB, CB, SL, JH), manuscript preparation (SB, SL, help from all authors), securing funding (JH) and project supervision (SL, JH).

## STAR Methods

### Resource availability

#### Lead Contact

Further information and requests for resources will be fulfilled upon reasonable request by the Lead Contact, Jochen Herms.

#### Materials Availability

This study did not generate new unique reagents or algorithms.

#### Data and Code Availability

Access to generated data and code will be made available upon reasonable request by the lead contact (JH).

#### Experimental Model and Subject Details

Wild-type C57BL/6 mice were obtained from the Jackson Laboratory. All animals were housed in groups under pathogen-free conditions and bred in the animal housing facility at the Center for Neuropathology and Prion Research of the Ludwig-Maximilians-University Munich, with food and water provided ad libitum (21 ± 2 °C, at 12/12 hour light/dark cycle). After cranial window implantation, mice were housed separately. All experiments were approved by the Bavarian government (Az. 55.2-1-54-2532-163-13) and performed according to the animal protection law.

## METHOD DETAILS

### PFF purification and seeding

Recombinant WT mouse a-synuclein was purified as previously described (Kostka et al., 2008; Nuscher et al., 2004). Preformed fibrils (PFFs) were assembled from purified a-synuclein monomer (5mg/ml) by incubation at 37°C and 1400 rpm for 96 hours and stored at −80°C (Conway et al., 2000). Directly before injection, an aliquot of PFFs was sonicated four times with a handheld probe (SonoPuls Mini 20, MS1.5, Bandelin, Berlin, Germany) according to the following protocol: Amplitude 30%; Time 15 s (pulse on 3s, pulse off 6 s). Two month-old mice were anesthetized with ketamine/xylazine (0.13/0.01 mg/g body weight; WDT/Bayer Health Care, Garbsen/Leverkusen, Germany) and stereotactically injected with 5 μl (25 μg) of PFFs into the dorsal striatum (coordinates relative to the Bregma: +0.2 mm anterior, +2.0 mm from midline, +2.6 mm beneath the dura) of the right hemisphere using a 5 μl Hamilton syringe. Injections were performed at 400 nl/min with the needle in place after injection for at least 5 min. Control animals received 5 μl sterile PBS. Animals were monitored regularly after the surgery.

### Virus injection and cranial window implantation

Eight months after PFF seeding, a cranial window was implanted over the right cortical hemisphere as previously reported (Holtmaat et al., 2009). In short, mice were given an intraperitoneal injection of ketamine/xylazine to reach surgical anesthesia. Additionally, dexamethasone (0.01 mg/g body weight; CP pharma, Burgdorf, Germany) was intraperitoneally administered immediately before surgery. A circular piece of the skull (4 mm in diameter) over the right hemisphere (approx. 1 mm caudal from the bregma and 3 mm lateral from midline) was removed, using a dental drill (Schick-Technikmaster C1; Pluradent; Offenbach, Germany).

AAV2/1.hSyn1.mRuby2.GSG.P2A.GCaMP6s.WPRE.SV4 was injected into 3-4 spots within the cranial window (300 nl per injection, 30 nl/min, 0.2 mm beneath the dura, virus titer ~ 10^12^ GC ml^−1^) The craniotomy was closed with a round coverslip (4 mm in diameter), held with dental acrylic (Hager & Werken, Duisburg, Germany). A small, L-shaped titan bar was glued next to the coverslip to allow repositioning of the mouse during subsequent imaging sessions. After surgery, mice received subcutaneous analgesic treatment with Carprophen (5 mg/kg body weight; Rimadyl; Pfizer, NY, USA) and antibiotic treatment with Cefotaxim (0.06 mg/kg body weight; Sanofi-Aventis, Frankfurt, Germany).

### *In vivo* imaging in awake, head-fixed mice

Imaging was performed in awake, head-fixed mice as previously reported (Guo et al., 2014). Mice were trained to head-fixation and the imaging equipment for at least 14 days prior to imaging. Mice were gradually habituated to the researcher, the holding tube and the setup noise. Head-fixation was applied for only several seconds at first, increasing with every training session up to 1 hour. Individual adjustments were taken if necessary to make the mice feel comfortable in their holding tube. During head-fixation, mice typically showed long episodes of quiet wakefulness (quiet) interrupted by brief episodes of intensive whisking and movement (active). Therefore, we used whisking movement as an indicator of behavioral state (quiet vs. active).

Imaging started 4 weeks after the cranial window preparation to allow the animals to recover from surgery and the window to clear. Before the imaging session in awake animals, mice were head-fixed and placed under the microscope for 5 min to habituate. *In vivo* imaging of neurons in layer 2/3 of somatosensory cortex (at a depth of 200 – 300 μm) was conducted using a La Vision Trim Scope (La Vision BioTec GmbH, Germany, 10 Hz frame rate, field of view (FOV) of 220 x 220 μm at 223 pixel resolution) equipped with a Ti:sapphire two-photon laser (Mai Tai, Spectra Physics, USA). Tuning of the laser to 940 nm enabled the simultaneous excitation of mRuby2 and GCaMP6s. Laser power was kept under 50 mW at all times, measured at the back-focal plane of the objective. Emitted fluorescence light was split at 560 nm and green light (495-560 nm, band pass filter) and red light (> 560nm) were detected by photomultiplier tubes. For recording mouse behavior, a web camera was installed and controlled by La Vison Imspector software which allowed synchronous recordings with same frame rate for mouse behavior and calcium transients. Activity of the same neurons was assessed under isoflurane anesthesia. To this end, mice were exposed to 0.5-1 vol % isoflurane over 10 min after which imaging commenced. Body temperature was kept stable with a heating pad and arterial saturation, breathing rate and heart rate were monitored with a pulse oxymeter (Starr Life Sciences).

### Data processing and image analysis

Collected images were processed and analyzed using custom-written codes in MATLAB and ImageJ software. Whisking was recorded by a camera (pco.pixelfly USB camera, Warner Instruments, Hamden, USA) synchronized with the imaging data. Based on whisker movement, the behavior in awake mice was classified as “active” or “quiet”. Two photon imaging data was registered to correct for slight brain displacement. Registration parameters (x/y shift) were estimated using a Fourier transform based approach (Guizar-Sicairos et al., 2008) on imaging data obtained from the mRuby2 (structural) channel (Rose et al., 2016). Regions of interest (ROIs) were manually identified using custom written software in MATLAB and a raw GCaMP6s fluorescence time series was constructed by averaging the pixel values within the ROI for each imaging frame. Time series were corrected for contamination by local neuropil fluorescence using the following method. Local neuropil fluorescence was calculated by averaging all pixel values in an approximately 30μm ring around the ROI, excluding other ROIs. The corrected ROI signal was computed based on the equation F_ROI_comp_ = F_ROI_ + 0.7x (median(F_neuropil_) − F _neuropil_) (Chen et al., 2013; Liebscher et al., 2016)).

F_ROI_comp_ represents the actual signal within the selected ROI after compensating for neuropil contamination, while F_ROI_ reflects the signal within the initially selected ROI. F_neuropil_ corresponds to the signal within the surrounding neuropil. Traces were next low pass filtered at 5Hz and slow fluctuations removed by subtracting the 8th percentile within a window of 1000 frames (equaling 100 seconds (Dombeck et al., 2007)). In order to estimate F0 we subtracted the 8th percentile in a very short sliding window of 1 second and used the median of all values below the 70th percentile of this ‘noise band’ as F0. ΔF/F then corresponds to F_ROI_comp_/F0. −1. ROIs were classified as active if ΔF/F exceeded 3x of the standard deviation of the noise band for at least 10 frames (1 second). For these active ROIs transient detection was performed on ΔF/F traces smoothed across 7 frames, using following criteria: >2 x standard deviation of the noise band minimum peak prominence of 0.15 and inter-peak interval minimum of 1 second. Whisking-associated neuronal activity was assessed by considering transients occurring within a window of 1 second before whisking onset up to 2 seconds after whisking offset. All other transients were considered spontaneous activity.

The correlation of neuronal activity for all pairs of active neurons in a given field of view was assessed by computing the Pearson correlation coefficient. To this end traces were smoothed over 25 frames and all values lower than 2x standard deviation of the noise band were set to 0. The correlation of activity during whisking and stationary epochs were analysed separately and compared to pairwise correlation derived from shuffled data. Shuffled data was generated by circularly shifting each trace at a random value between 1 to the length of the activity trace. For each field of view the average of all pairwise correlations was computed and those values were compared between control and a-syn mice.

In total we recorded from 1561 neurons in 32 experiments (field of views) from 7 control mice and 1534 neurons in 32 experiments from 7 a-syn mice.

### Immunohistochemistry

After imaging, mice (control, n = 5, a-syn, n = 5) were sacrificed by transcardial perfusion with phosphate-buffered saline (PBS) followed by 4% paraformaldehyde (w/v) in deep ketamine/xylazine anesthesia. The brains were removed and post-fixed in PBS containing 4% paraformaldehyde over night before cutting 50μm thick coronal sections on a vibratome (VT 1000 S from Leica, Wetzlar, Germany). Floating sections were permeabilized with 2% Triton X-100 in PBS overnight and blocked with 2% normal goat serum (Sigma-Aldrich) and 4% BSA (Bovine Serum Albumin, VWR) for 4h on a shaker at room temperature. Primary antibodies (Anti-Alpha-synuclein - phospho S129, rabbit polyclonal, Abcam, Cambridge, UK; Anti-GAD67, mouse monoclonal, Millipore, CA, USA) were incubated for 48h at 4°C in a 1:1000 dilution, according to the manufacturer’s recommendations. Slices were washed 3 × 10min with PBS and then incubated with the secondary antibodies (1:1000; goat anti-rabbit Alexa 488, goat antimouse Alexa 647, Invitrogen, Life Technologies GmbH) for overnight at 4°C. After 3 × 10min washing in PBS, sections were incubated for 1h with NeuroTrace 530/615 Red Fluorescent Nissl Stain (1:500, ThermoFisher Scientific) on a shaker at room temperature. Sections were finally washed for 3 × 10min with PBS before mounting them on glass coverslips with Dako fluorescence conserving media (Dako, Hamburg, Germany).

### Stereology

Brain sections outlines were carried out using a Zeiss fluorescent microscope (Imager.M2, ZEISS, Oberkochen, Germany) fully motorized and interfacing to a Dell computer running StereoInvestigator^®^ (MBF Bioscience, Williston, Vermont, USA). Cells labeled with corresponding markers were quantified in 7 serial coronal sections spanning the entire brain hemisphere in the coronal plain, spaced 600μm apart respectively. Outline and fiduciary marks were drawn at 2.5x magnification (EC-Plan-NEOFLUAR 2.5X/0.075, ZEISS, Oberkochen, Germany) using Neurotrace stain to delineate reference points. Limits for areas of interest were drawn following a mouse brain Atlas (Paxinos and Franklin, 2013). Cells were identified as positive for a marker if they expressed immunoreactivity visually deemed to be above background even if it was very weak, which means cells exhibiting varying levels of immunolabeling from very weakly to very strongly stained were all identified as marker positive. All cell counting was done by an investigator blind to genotype and treatment, at 63x magnification (Plan/APOCHROMAT 63X/1.4 Oil DIC, ZEISS, Oberkochen, Germany) using a 3D counting frame in a sampling grid (Suppl. Table 1). The coefficient of error (Gundersen), m=1, and the average cell counts per sampling site are described for each marker and region (Suppl. Table 1).

### Statistics

If not stated otherwise, we employed student’s t-test to compare the average of normally distributed data (e.g. population response to whisking). For non-normally distributed data we used the ranksum (Mann-Whitney-U) test (e.g. fraction of whisking responsive neurons). Distribution of data was compared using the Kolmogorov-Smirnov test (KS, e.g. distribution of frequencies, amplitudes, correlation coefficients). Stereology results were compared using a two-way ANOVA followed by Bonferroni’s *post hoc* test.

---

**Suppl. Table 1.** Parameters for stereological cell counting in the cortex as well as the coefficient error and the average cell count per sampling site for each marker

| Ctx SSp |  | Counting frame | Sampling<br>grid size | Coefficient of<br>error (Gundersen),<br>m=1 | Average cell counts<br>/sampling site |
| --- | --- | --- | --- | --- | --- |
| GAD67 | PBS | 100x100x8 $\mu\text{m}$ | 300x300 $\mu\text{m}$ | 0.12 | 1.10 |
| | PFF | 100x100x8 $\mu\text{m}$ | 300x300 $\mu\text{m}$ | 0.13 | 0.84 |
| Neurotrace | PBS | 20 x 20 x12 $\mu\text{m}$ | 200x200 $\mu\text{m}$ | 0.11 | 1.22 |
| | PFF | 20 x 20 x12 $\mu\text{m}$ | 200x200 $\mu\text{m}$ | 0.11 | 1.14 |

**Suppl. Fig. S1:**
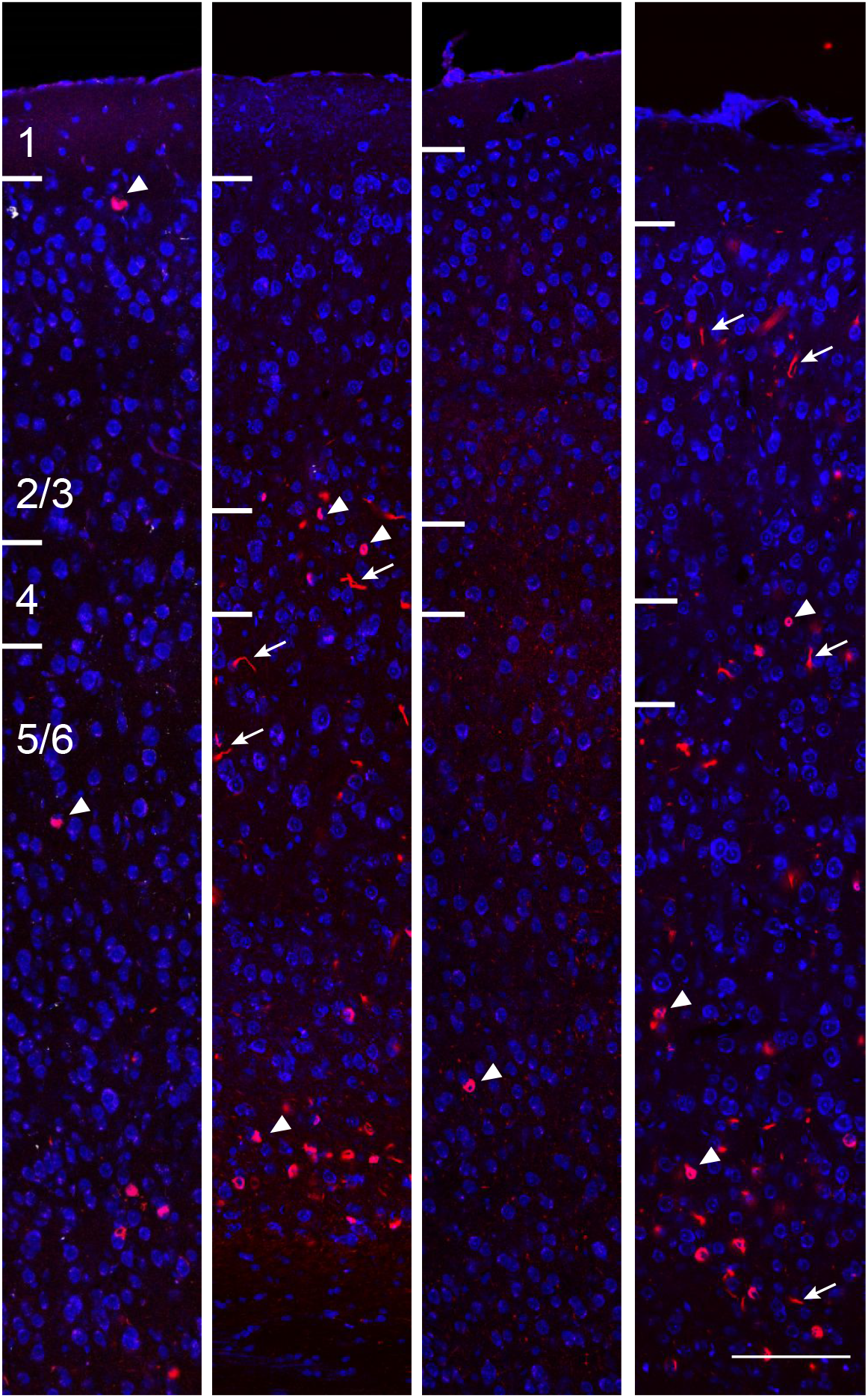
Striatal a-syn PFF seeding causes the formation of Lewy-body like aggregates in cortex of wild-type mice. Examples of neuritic (arrows) and somatic (arrowheads) a-syn aggregates (red) across all cortical layers in S1. (blue – neurotrace), Scale bar 100μm.

**Suppl. Figure S2.**
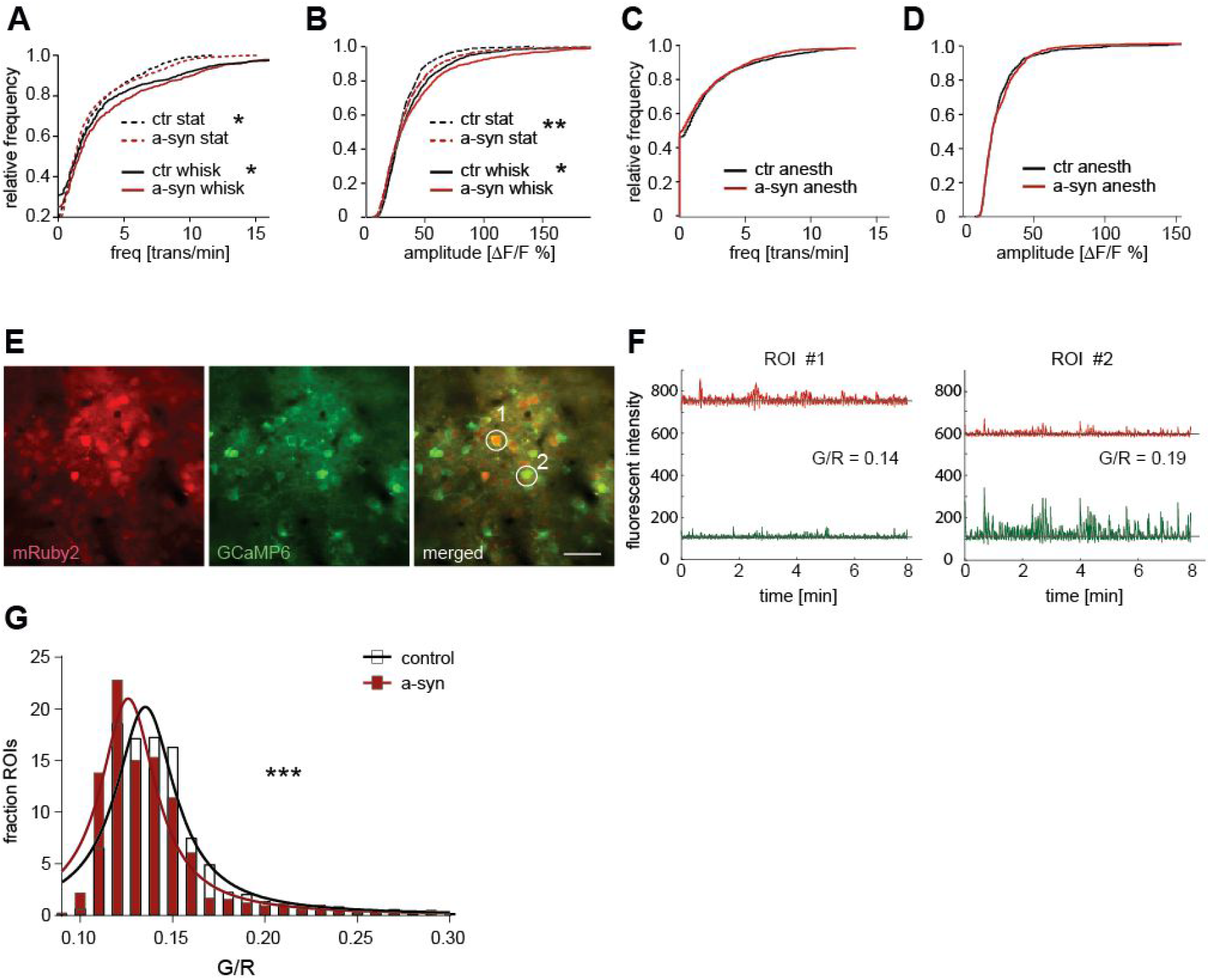
Neuronal activity in S1 is altered upon striatal seeding of a-syn PFF. **(A)** Neuronal activity in S1 is increased in a-syn mice both during quiescence as well as during whisking-associated epochs. **(B)** Transient amplitudes during quiescence and during whisking-associated epochs are larger in a-syn mice. **(C)** Spontaneous neuronal frequency and **(D)** transient amplitudes did not differ between control and a-syn mice during anesthesia. (E) Both channels of a representative field of view are shown (average projection of all frames), highlighting the difference in expression of mRuby2 and GCaMP6s in individual cells. (F) Fluorescent signal intensities of 2 ROIs (marked in (E)) of both channels are depicted together with the corresponding green/red ratio (G/R), based on the baseline values obtained for each ROI in the red and green channel (marked by thin black line superimposed on the trace). (G) Distribution of G/R ratios for all ROIs overlaid with a Lorentzian fit. Scale bar in (E) 50 μm, * *P* < 0.05, ** *P* < 0.001. *** *P* < 0.0001.

## References

Arnoux, I., Willam, M., Griesche, N., Krummeich, J., Watari, H., Offermann, N., Weber, S., Narayan Dey, P., Chen, C., Monteiro, O., et al. (2018). Metformin reverses early cortical network dysfunction and behavior changes in Huntington’s disease. eLife 7, e38744.

Betzer, C., Lassen, L.B., Olsen, A., Kofoed, R.H., Reimer, L., Gregersen, E., Zheng, J., Calì, T., Gai, W.P., Chen, T., et al. (2018). Alpha-synuclein aggregates activate calcium pump SERCA leading to calcium dysregulation. EMBO reports 19, e44617.

Blumenstock, S., Rodrigues, E.F., Peters, F., Blazquez-Llorca, L., Schmidt, F., Giese, A., and Herms, J. (2017). Seeding and transgenic overexpression of alpha-synuclein triggers dendritic spine pathology in the neocortex. EMBO Molecular Medicine 9, 716–731.

Burgold, J., Schulz-Trieglaff, E.K., Voelkl, K., Gutiérrez-Ángel, S., Bader, J.M., Hosp, F., Mann, M., Arzberger, T., Klein, R., Liebscher, S., et al. (2019). Cortical circuit alterations precede motor impairments in Huntington’s disease mice. Scientific Reports 9, 6634.

Busche, M.A., Eichhoff, G., Adelsberger, H., Abramowski, D., Wiederhold, K.-H., Haass, C., Staufenbiel, M., Konnerth, A., and Garaschuk, O. (2008). Clusters of Hyperactive Neurons Near Amyloid Plaques in a Mouse Model of Alzheimer’s Disease. Science 321, 1686–1689.

Challis, C., Hori, A., Sampson, T.R., Yoo, B.B., Challis, R.C., Hamilton, A.M., Mazmanian, S.K., Volpicelli-Daley, L.A., and Gradinaru, V. (2020). Gut-seeded α-synuclein fibrils promote gut dysfunction and brain pathology specifically in aged mice. Nature Neuroscience 23, 327–336.

Chen, Q.-Q., Haikal, C., Li, W., Li, M.-T., Wang, Z.-Y., and Li, J.-Y. (2018). Age-dependent alpha-synuclein accumulation and aggregation in the colon of a transgenic mouse model of Parkinson’s disease. Translational Neurodegeneration 7, 13.

Chen, T.-W., Wardill, T.J., Sun, Y., Pulver, S.R., Renninger, S.L., Baohan, A., Schreiter, E.R., Kerr, R.A., Orger, M.B., Jayaraman, V., et al. (2013). Ultrasensitive fluorescent proteins for imaging neuronal activity. Nature 499, 295–300.

Conway, K.A., Harper, J.D., and Lansbury, P.T. (2000). Fibrils Formed in Vitro from α-Synuclein and Two Mutant Forms Linked to Parkinson’s Disease are Typical Amyloid \textsuperscript†. Biochemistry 39, 2552–2563.

Demuro, A., Mina, E., Kayed, R., Milton, S.C., Parker, I., and Glabe, C.G. (2005). Calcium Dysregulation and Membrane Disruption as a Ubiquitous Neurotoxic Mechanism of Soluble Amyloid Oligomers. Journal of Biological Chemistry 280, 17294–17300.

Dombeck, D.A., Khabbaz, A.N., Collman, F., Adelman, T.L., and Tank, D.W. (2007). Imaging large-scale neural activity with cellular resolution in awake, mobile mice. Neuron 56, 43–57.

Guizar-Sicairos, M., Thurman, S.T., and Fienup, J.R. (2008). Efficient subpixel image registration algorithms. Optics letters 33, 156–158.

Guo, Z.V., Hires, S.A., Li, N., O’Connor, D.H., Komiyama, T., Ophir, E., Huber, D., Bonardi, C., Morandell, K., Gutnisky, D., et al. (2014). Procedures for behavioral experiments in head-fixed mice. PloS One 9, e88678.

Henderson, M.X., Cornblath, E.J., Darwich, A., Zhang, B., Brown, H., Gathagan, R.J., Sandler, R.M., Bassett, D.S., Trojanowski, J.Q., and Lee, V.M.Y. (2019). Spread of α-synuclein pathology through the brain connectome is modulated by selective vulnerability and predicted by network analysis. Nature Neuroscience 22, 1248–1257.

Hettiarachchi, N.T., Parker, A., Dallas, M.L., Pennington, K., Hung, C.-C., Pearson, H.A., Boyle, J.P., Robinson, P., and Peers, C. (2009). α-Synuclein modulation of Ca2+ signaling in human neuroblastoma (SH-SY5Y) cells. Journal of Neurochemistry 111, 1192–1201.

Holtmaat, A., Bonhoeffer, T., Chow, D.K., Chuckowree, J., De Paola, V., Hofer, S.B., Hübener, M., Keck, T., Knott, G., Lee, W.-C.A., et al. (2009). Long-term, high-resolution imaging in the mouse neocortex through a chronic cranial window. Nature Protocols 4, 1128–1144.

Hüls, S., Högen, T., Vassallo, N., Danzer, K.M., Hengerer, B., Giese, A., and Herms, J. (2011). AMPA-receptor-mediated excitatory synaptic transmission is enhanced by iron-induced α-synuclein oligomers: α-Synuclein oligomers alter synaptic transmission. Journal of Neurochemistry 117, 868–878.

Kostka, M., Högen, T., Danzer, K.M., Levin, J., Habeck, M., Wirth, A., Wagner, R., Glabe, C.G., Finger, S., Heinzelmann, U., et al. (2008). Single Particle Characterization of Iron-induced Pore-forming α-Synuclein Oligomers. Journal of Biological Chemistry 283, 10992–11003.

Kulkarni, A.S., del Mar Cortijo, M., Roberts, E.R., Suggs, T.L., Stover, H.B., Pena-Bravo, J.I., Steiner, J.A., Luk, K.C., Brundin, P., and Wesson, D.W. (2020). Perturbation of in vivo neural activity following α-Synuclein seeding in the olfactory bulb. (Neuroscience).

Li, J.-Y., Englund, E., Holton, J.L., Soulet, D., Hagell, P., Lees, A.J., Lashley, T., Quinn, N.P., Rehncrona, S., Björklund, A., et al. (2008). Lewy bodies in grafted neurons in subjects with Parkinson’s disease suggest host-to-graft disease propagation. Nature Medicine 14, 501–503.

Li, Q., Ke, Y., Chan, D.C.W., Qian, Z.-M., Yung, K.K.L., Ko, H., Arbuthnott, G.W., and Yung, W.-H. (2012). Therapeutic Deep Brain Stimulation in Parkinsonian Rats Directly Influences Motor Cortex. Neuron 76, 1030–1041.

Liebscher, S., Keller, G.B., Goltstein, P.M., Bonhoeffer, T., and Hübener, M. (2016). Selective Persistence of Sensorimotor Mismatch Signals in Visual Cortex of Behaving Alzheimer’s Disease Mice. Current Biology 26, 956–964.

Luk, K.C., Kehm, V., Carroll, J., Zhang, B., O’Brien, P., Trojanowski, J.Q., and Lee, V.M.-Y. (2012). Pathological α-Synuclein Transmission Initiates Parkinson-like Neurodegeneration in Nontransgenic Mice. Science 338, 949–953.

Marinković, P., Blumenstock, S., Goltstein, P.M., Korzhova, V., Peters, F., Knebl, A., and Herms, J. (2019). In vivo imaging reveals reduced activity of neuronal circuits in a mouse tauopathy model. Brain 142, 1051–1062.

Masuda-Suzukake, M., Nonaka, T., Hosokawa, M., Oikawa, T., Arai, T., Akiyama, H., Mann, D.M.A., and Hasegawa, M. (2013). Prion-like spreading of pathological α-synuclein in brain. Brain 136, 1128–1138.

McGregor, M.M., and Nelson, A.B. (2019). Circuit Mechanisms of Parkinson’s Disease. Neuron 101, 1042–1056.

Nagahama, Y., Okina, T., Suzuki, N., and Matsuda, M. (2010). Neural correlates of psychotic symptoms in dementia with Lewy bodies. Brain 133, 557–567.

Nuscher, B., Kamp, F., Mehnert, T., Odoy, S., Haass, C., Kahle, P.J., and Beyer, K. (2004). α-Synuclein Has a High Affinity for Packing Defects in a Bilayer Membrane A THERMODYNAMICS STUDY. Journal of Biological Chemistry 279, 21966–21975.

Osterberg, V.R., Spinelli, K.J., Weston, L.J., Luk, K.C., Woltjer, R.L., and Unni, V.K. (2015). Progressive Aggregation of Alpha-Synuclein and Selective Degeneration of Lewy Inclusion-Bearing Neurons in a Mouse Model of Parkinsonism. Cell Reports 10, 1252–1260.

Parker, J.G., Marshall, J.D., Ahanonu, B., Wu, Y.-W., Kim, T.H., Grewe, B.F., Zhang, Y., Li, J.Z., Ding, J.B., Ehlers, M.D., et al. (2018). Diametric neural ensemble dynamics in parkinsonian and dyskinetic states. Nature.

Paxinos, G., and Franklin, K.B. (2013). Paxinos and Franklin’s the Mouse Brain in Stereotaxic Coordinates, Compact - 5th Edition.

Peelaerts, W., Bousset, L., Van der Perren, A., Moskalyuk, A., Pulizzi, R., Giugliano, M., Van den Haute, C., Melki, R., and Baekelandt, V. (2015). α-Synuclein strains cause distinct synucleinopathies after local and systemic administration. Nature 522, 340–344.

Peng, C., Trojanowski, J.Q., and Lee, V.M.-Y. (2020). Protein transmission in neurodegenerative disease. Nature Reviews Neurology 16, 199–212.

Reznichenko, L., Cheng, Q., Nizar, K., Gratiy, S.L., Saisan, P.A., Rockenstein, E.M., González, T., Patrick, C., Spencer, B., Desplats, P., et al. (2012). In Vivo Alterations in Calcium Buffering Capacity in Transgenic Mouse Model of Synucleinopathy. The Journal of Neuroscience 32, 9992–9998.

Rose, T., Jaepel, J., Hübener, M., and Bonhoeffer, T. (2016). Cell-specific restoration of stimulus preference after monocular deprivation in the visual cortex. Science (New York, N.Y.) 352, 1319–1322.

Schmidt, F., Levin, J., Kamp, F., Kretzschmar, H., Giese, A., and Bötzel, K. (2012). Single-Channel Electrophysiology Reveals a Distinct and Uniform Pore Complex Formed by α-Synuclein Oligomers in Lipid Membranes. PLoS ONE 7, e42545.

Shrivastava, A.N., Aperia, A., Melki, R., and Triller, A. (2017). Physico-Pathologic Mechanisms Involved in Neurodegeneration: Misfolded Protein-Plasma Membrane Interactions. Neuron 95, 33–50.

Shrivastava, A.N., Redeker, V., Fritz, N., Pieri, L., Almeida, L.G., Spolidoro, M., Liebmann, T., Bousset, L., Renner, M., Lena, C., et al. (2015). α-synuclein assemblies sequester neuronal 3-Na+/K+-ATPase and impair Na+ gradient. The EMBO Journal 34, 2408–2423.

Spillantini, M.G., Crowther, R.A., Jakes, R., Hasegawa, M., and Goedert, M. (1998). α-Synuclein in filamentous inclusions of Lewy bodies from Parkinson’s disease and dementia with Lewy bodies. Proceedings of the National Academy of Sciences 95, 6469–6473.

Spillantini, M.G., Schmidt, M.L., Lee, V.M.-Y., Trojanowski, J.Q., Jakes, R., and Goedert, M. (1997). α-Synuclein in Lewy bodies. Nature 388, 839–840.

Uchihara, T., and Giasson, B.I. (2016). Propagation of alpha-synuclein pathology: hypotheses, discoveries, and yet unresolved questions from experimental and human brain studies. Acta Neuropathologica 131, 49–73.

Volles, M.J., Lee, S.-J., Rochet, J.-C., Shtilerman, M.D., Ding, T.T., Kessler, J.C., and Lansbury, P.T. (2001). Vesicle Permeabilization by Protofibrillar α-Synuclein: Implications for the Pathogenesis and Treatment of Parkinson’s Disease. Biochemistry 40, 7812–7819.

Volpicelli-Daley, L.A., Luk, K.C., Patel, T.P., Tanik, S.A., Riddle, D.M., Stieber, A., Meaney, D.F., Trojanowski, J.Q., and Lee, V.M.-Y. (2011). Exogenous α-Synuclein Fibrils Induce Lewy Body Pathology Leading to Synaptic Dysfunction and Neuron Death. Neuron 72, 57–71.

Wang, X.-J., Ma, M.-M., Zhou, L.-B., Jiang, X.-Y., Hao, M.-M., Teng, R.K.F., Wu, E., Tang, B.-S., Li, J.-Y., Teng, J.-F., et al. (2020). Autonomic ganglionic injection of α-synuclein fibrils as a model of pure autonomic failure α-synucleinopathy. Nature Communications 11, 934.

Wong, Y.C., and Krainc, D. (2017). α-synuclein toxicity in neurodegeneration: mechanism and therapeutic strategies. Nature Medicine 23, 1–13.

Wu, Q., Takano, H., Riddle, D.M., Trojanowski, J.Q., Coulter, D.A., and Lee, V.M. (2019). alpha-Synuclein (alphaSyn) Preformed Fibrils Induce Endogenous alphaSyn Aggregation, Compromise Synaptic Activity and Enhance Synapse Loss in Cultured Excitatory Hippocampal Neurons. J Neurosci 39, 5080–5094.

Yu, H., Sternad, D., Corcos, D.M., and Vaillancourt, D.E. (2007). Role of hyperactive cerebellum and motor cortex in Parkinson’s disease. NeuroImage 35, 222–233.

